# A Cell-Permeable Nanobody to Restore F508del Cystic Fibrosis Transmembrane Conductance Regulator Activity

**DOI:** 10.1101/2024.04.26.591242

**Authors:** Luise Franz, Tihomir Rubil, Anita Balázs, Marie Overtus, Kristin Kemnitz-Hassanin, Cedric Govaerts, Marcus A. Mall, Christian P.R. Hackenberger

## Abstract

Nanobodies have gained considerable attention as particularly promising biopharmaceuticals. However, nanobody-based modalities are currently limited to extracellular targets due to a lack of efficient delivery methods required to reach targets inside cells. In this study, we introduce cell-permeable nanobodies for targeting a disease-relevant intracellular protein, namely the cystic fibrosis transmembrane conductance regulator (CFTR) chloride channel with the most common cystic fibrosis (CF)-causing mutation F508del. We employ cell-penetrating peptides (CPPs) to deliver a CFTR-binding nanobody (NB1) that stabilizes misfolded F508del-CFTR and prevents its degradation to restore its function. Our data show that conjugation of a disulfide-linked CPP in combination with a cell-surface anchored CPP-additive enables intracellular delivery of NB1 into CF bronchial epithelial cells, which promotes maturation and trafficking of F508del-CFTR protein to the apical cell membrane. Furthermore, we demonstrate that the cell-permeable nanobody restores CFTR chloride channel function, which can be further enhanced by the clinically approved small molecule CFTR potentiator ivacaftor. This study highlights the use of cell-permeable nanobodies for modulation of protein function and illustrates their therapeutic potential as next-generation biopharmaceuticals for intracellular delivery and targeting.

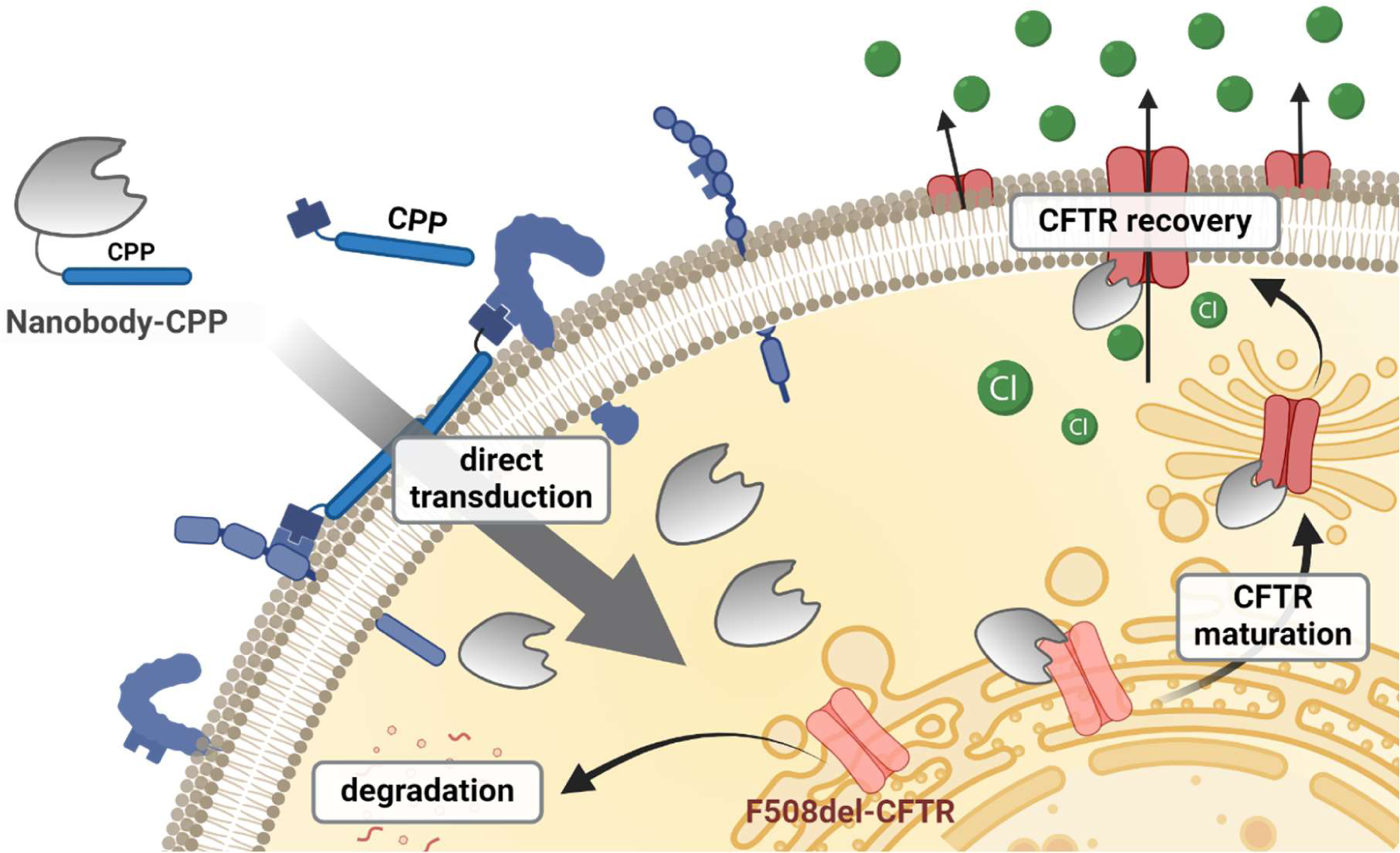

## Main

Biopharmaceuticals are on the rise in pharmaceutical development, with monoclonal antibodies making up over half of the approved biopharmaceuticals in the US and Europe between 2018-2022^1^. While they are widely applied in treatment of cancer, inflammation, and metabolic diseases, their application is limited by their inability for deep tissue penetration due to their size and immunogenicity^2^. Antibody fragments, e.g. single chain variable fragments, and nanobodies, derived from the variable domain of heavy chain only antibodies^3^, have received much attention to overcome these limitations. In particular, nanobodies pose several attractive features including high solubility, high temperature and physicochemical stability^4, 5^ and long shelf live.^6^ Large synthetic libraries for the selection and development of new targeted binders are available.^7, 8^ Moreover, nanobodies are straightforwardly accessible by expression in bacteria and can easily be modified by enzymatic or chemical modification strategies.^4, 7^ These aspects resulted in a wide array of applications in the biological and pharmacological sciences, which are further facilitated by their fast clearance, deep tissue penetration and low immunogenicity compared to conventional antibodies^9^.

Despite great advances in developing nanobodies for therapeutic applications, current nanobody modalities are limited to address extracellular targets due to a lack of cell-permeable variants. Nevertheless, it is well established that nanobodies are able to address intracellular targets to manipulate, modulate, inhibit or degrade relevant proteins in form of intracellularly expressed nanobodies^7^. These so-called intrabodies rely on the delivery of genetic material into the cell to express the nanobody, which is suitable *in cellulo* but necessitates gene therapy in a clinical context. Additionally, intrabodies do not allow for non-genetic modification, which prevents enzymatic or chemical modifications for labeling or stabilization.

In recent years, several approaches have been pursued for the intracellular delivery of proteins and nanobodies, including physical methods to overcome the cell membrane by pore formation or membrane disruption^10–13^, as well as injection using a bacterial type III protein secretion system as a “molecular syringe.”^14, 15^ Furthermore, nanobody supercharging^16^ and the use of cyclic arginine-rich cell-penetrating peptides (CPPs)^17^ or cell surface anchoring CPP-additives have been used^18^. However, proof-of-concept studies with cell-permeable nanobodies so far focused mainly on targeting GFP in engineered cell lines or on the labeling of endogenous protein targets, in particular for super resolution microscopy.^19^

To demonstrate the prospect of cell-permeable nanobodies to modulate a therapeutically highly relevant intracellular target, we now present a cell-permeable nanobody, which reconstitutes the activity of the misfolded cystic fibrosis transmembrane conductance regulator (CFTR) causing cystic fibrosis (CF). The most common mutation in CF patients is a deletion of F508 in CFTR, which causes misfolding and intracellular degradation of CFTR anion channels, thus incapacitating insertion into the apical membrane, resulting in impaired transepithelial transport of chloride and bicarbonate, which are essential for host defense and homeostasis in the lungs and other epithelial organs^20, 21^. Current therapeutic approaches focus on pharmacologic rescue of F508del-CFTR using a combination of the small molecule CFTR correctors elexacaftor and tezacaftor in combination with the potentiator ivacaftor^22, 23^. While this triple combination therapy provides unprecedented improvement in clinical outcomes of patients with at least one copy of the common F508del mutation^24^, restoration of CFTR function remains partial, and chronic infection and inflammation of the lungs persist,^25–28^ underscoring the need for further optimization of F508del correction^20^.

Recently, a CFTR-binding nanobody was shown to stabilize F508del-CFTR *in vitro* suggesting the potential to correct the folding defect and restore CFTR function^29^. However, therapeutic development of CFTR-targeting nanobodies has been hampered by the lack of tools for intracellular delivery, where CPPs could offer a technological advance to bridge this gap. Here, we selected a previously reported nanobody^29^ that exhibited high affinity against wild-type and F508del-CFTR to demonstrate the utility of cell-permeable nanobodies as tools to intracellularly modulate endogenous protein functionality. To render the nanobody cell-permeable, we proposed to use our previously introduced cell-penetrating peptide (CPP)-additive strategy, which enabled the cytosolic delivery of protein-CPP conjugates at low concentrations^18, 19^. We investigated the effects of intracellular nanobody delivery on maturation, trafficking and chloride channel function of F508del-CFTR utilizing a variety of biochemical methods including live-cell fluorescence microscopy, flow cytometry, Western blot analysis, and transepithelial short-circuit current measurements.

## Results

### Design of a cell permeable F508del-CFTR targeting nanobody

We started our investigation by expressing a suitable cysteine containing nanobody variant of a reported CFTR binding nanobody^29^ further referred to as NB1, to allow the attachment of a CPP (R_10_)-peptide via a disulfide. Thereby, the NB1-CPP conjugate can be taken up by the cell by direct transduction and the disulfide conjugated CPP can be cleaved in the intracellular reductive environment to ensure intracellular target engagement of the nanobody^17, 30^. In addition, NB1 contains an *N*-terminal glycine after processing for sortase-mediated site-specific fluorophore labeling to track successful cellular uptake (Fig. 1A). Since the incorporation of a *C*-terminal cysteine can drastically reduce the yield of bacterial nanobody expression^31^, we optimized the expression and purification of NB1. Expression was performed in TB-medium with additional glucose and Mg_2_Cl, to increase the protein yield. Furthermore, reductive conditions during all purification steps ensured a reduction of loss of protein due to dimerization. The changes resulted in an overall 6-fold increase in yield reaching 3 mg/L expression culture in high purity (Fig. 1B, i and ii) compared to the previously established protocol for the unaltered nanobody variant^29^. *N-* terminal sortase labeling^32^ with 1.1 equivalents of sortase and 20 eq. of Fluorescein isothiocyanate (FITC)-LPETGG peptide and subsequent sortase removal yielded the desired fluorescent nanobody cys-mutant (FITC-NB1). The final CPP conjugation was performed as previously described^18^ with 3 eq. of TNB-R_10_ overnight at 4°C to yield the cell-permeable fluorescent CPP-disulfide-conjugated nanobody (FITC-NB1-R_10_), in which we monitored labeling and conjugation success by SDS-PAGE (Fig. 1C).

**Figure 1.**
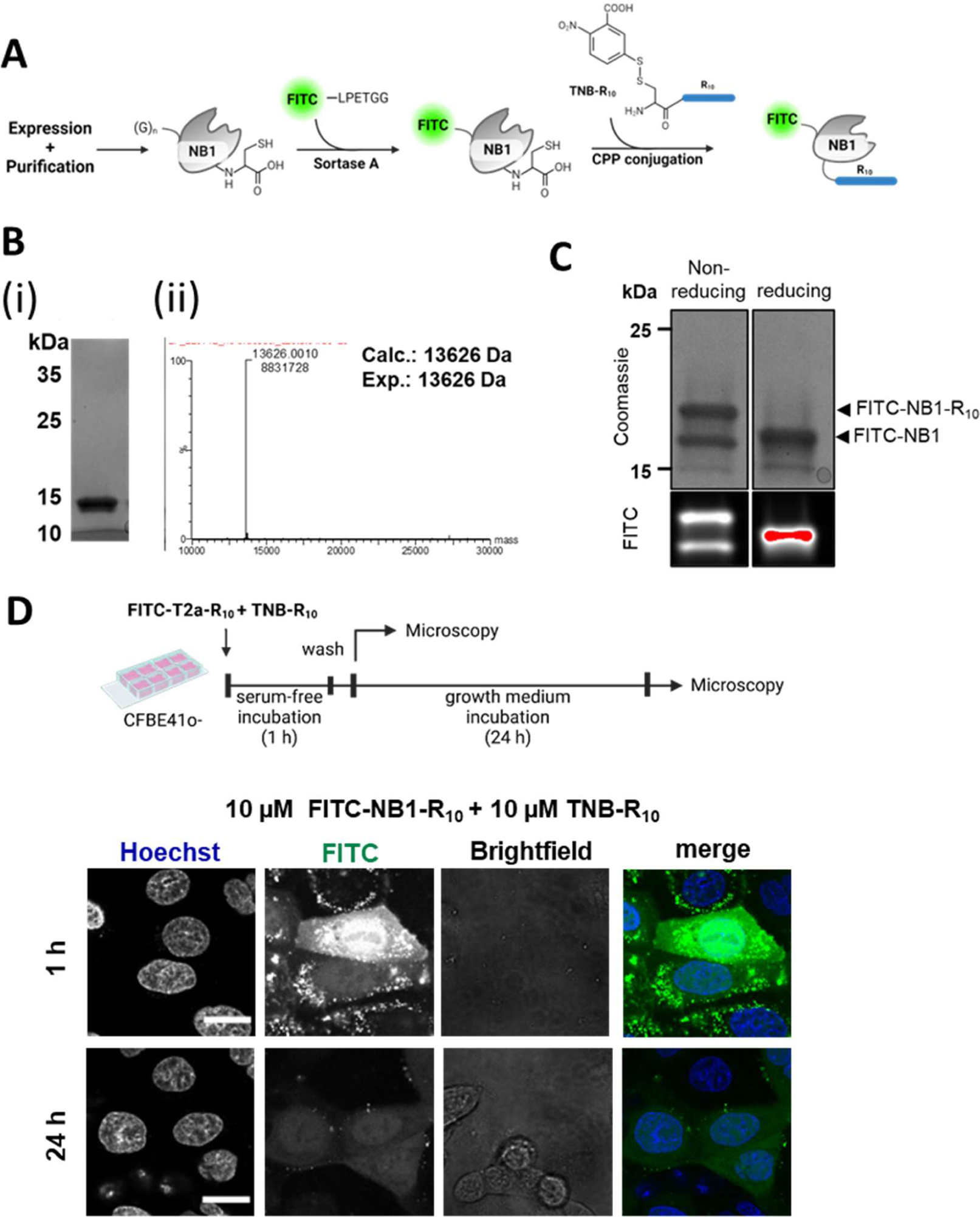
Design and intracellular delivery of a CFTR-binding nanobody. **A** CPP-conjugation and fluorescent labeling scheme. **B** Coomassie stained SDS-PAGE of NB1 (i) after purification and (ii) deconvoluted HR-MS analysis of expressed and purified NB1 nanobody Cys-mutant **C** SDS-PAGE analysis of FITC labeled and CPP-conjugated NB1 (FITC-NB1-R10) treated with 2.5% β-mercaptoethanol (reducing) or without (non-reducing) in Laemmli buffer. Fluorescence image (FITC) taken prior to Coomassie staining (Coomassie). **D** Live-cell microscopy images of cellular delivery in CFBE41o-cells with 10 µM FITC-NB1-R10/10 µM TNB-R10 directly after 1 h of incubation in serum-free medium (1h) or after subsequent 24 h of incubation in growth medium (24 h). Scale bar: 20 µM.

### Determining ideal treatment conditions of cell-permeable nanobody for F508del-CFTR recovery

For the delivery of the fluorescent CPP-nanobody-conjugate FITC-NB1-R_10_ we used the thiol-reactive polyarginine-containing CPP (TNB-R_10_), used prior for CPP conjugation, as an additive^18^. First, we probed the delivery of FITC-NB1-R_10_ into different human cell lines (HeLa, HEK, A549). We observed successful cytosolic delivery in all cell lines at 10 µM FITC-NB1-R_10_/10 µM TNB-R_10_, determined by diffuse fluorescence signal across the cytosol and nucleus of the cell (Extended Data Fig. 1)^33^. Nevertheless, the uptake efficiency varied between the cell lines with uptake being observed at concentrations as low as 2.5 µM FITC-NB1-R_10_/10 µM TNB-R_10_ in HeLa and HEK cells, while successful cytosolic uptake in A549 cells was exclusively observed at 10 µM FITC-NB1-R_10_/10 µM TNB-R_10_.

Next, we tested the uptake in CF bronchial epithelial cells (CFBE41o-) expressing F508del-CFTR, a disease-relevant model of the CF airway epithelium suitable for functional studies^34 35^. In CFBE41o-cells, ideal uptake was first observed at 10 µM FITC-NB1-R_10_/10 µM TNB-R_10_ (Fig. 1D), indicated by the clear fluorescence signal across the cytosol and the nucleus. Importantly, even after an additional 24 h incubation in growth medium, a fluorescent signal, although weaker, could be detected in the cytosol of cells indicating the persistence of the nanobody in the cell over that time (Fig. 1D).

Based on these results, we investigated if a non-fluorescent CPP-nanobody conjugate (NB1-R_10_) could intracellularly stabilize F508del and restore CFTR-mediated chloride transport in CFBE41o-cells after cellular delivery. To that end, we performed transepithelial short-circuit current (I_sc_) measurements in Ussing chambers as a dose-response study and to determine functional uptake conditions. CFBE41o-cells were cultured as a confluent monolayer on semi-permeable filters and incubated apically for 24 h with NB1-R_10_ and TNB-R_10_ additives in serum-free medium. CFTR function was determined by measuring changes in I_sc_ in response to cAMP-dependent activation with forskolin/IBMX, and inhibition with CFTR inhibitor-172 (CFTRinh-172). Initial measurements revealed that incubation with 10 µM NB1-R_10_/10 µM TNB-R_10_, while suitable in uptake experiments, did not restore CFTR function in Ussing chamber experiments (Extended Data Fig. 2A, B). Further dose-response studies revealed that a concentration of at least 50 µM NB1-R_10_ was necessary to rescue CFTR function in CFBE41o-monolayers, as evidenced by substantial increases in forskolin/IBMX-induced and CFTRinh-172-sensitive I_sc_. An optimal effect was observed at 75 µM NB1-R_10_/30 µM TNB-R_10_ and increasing the concentrations of both nanobody and additive beyond these levels did not lead to any additional improvements in CFTR function (Extended Data Fig. 2A, B). Subsequent uptake and cytotoxicity experiments at these conditions confirmed that the chosen nanobody concentrations are not toxic for the cells (Extended Data Fig. 2C). Uptake experiments under these conditions revealed a strong signal after 24 h in cells (Extended Data Fig. 2D) indicating that the higher amount of nanobody might be necessary to allow for sufficient intracellular concentrations of NB1 to achieve the restoration of function of mutated CFTR.

### Nanobody treatment rescues F508del maturation, trafficking and function in CF bronchial epithelial cells

After determining ideal treatment conditions that affect CFTR function upon cellular uptake of NB1-R_10_, we further investigated the effect of the cell-permeable CFTR-nanobody NB1-R_10_ on F508del-CFTR maturation and trafficking, which could lead to the functional recovery of F508del-CFTR. Therefore, we first evaluated CFTR-maturation upon cell-permeable nanobody treatment by Western blot analysis of CFBE41o-cell lysates under previously determined ideal conditions (75 µM TNB-R_10_/30 µM TNB-R_10_). Thus, cells were cultured as confluent monolayers on semi-permeable filters and treated apically with 75 µM TNB-R_10_/30 µM TNB-R_10_ for 24 h in serum-free medium. As positive control, CFBE41o-cells were incubated at low temperature (27 °C) for 24 h, which was previously shown to restore folding and function of F508del-CFTR^36^. We observed two specific bands in both nanobody-treated and temperature-corrected (27 °C) groups that correspond to the core glycosylated (B-band) and complex glycosylated (C-band) form of CFTR (Fig. 2A), whereas the untreated cell lysates showed only the presence of the B-band^22, 36^. Densitometric analysis confirmed that the relative amount of CFTR C-band protein was significantly increased in both NB1-R_10_ and 27°C groups compared to no treatment controls (Fig. 2B), demonstrating increased CFTR maturation associated with increased trafficking from the endoplasmic reticulum to the Golgi apparatus.

**Figure 2.**
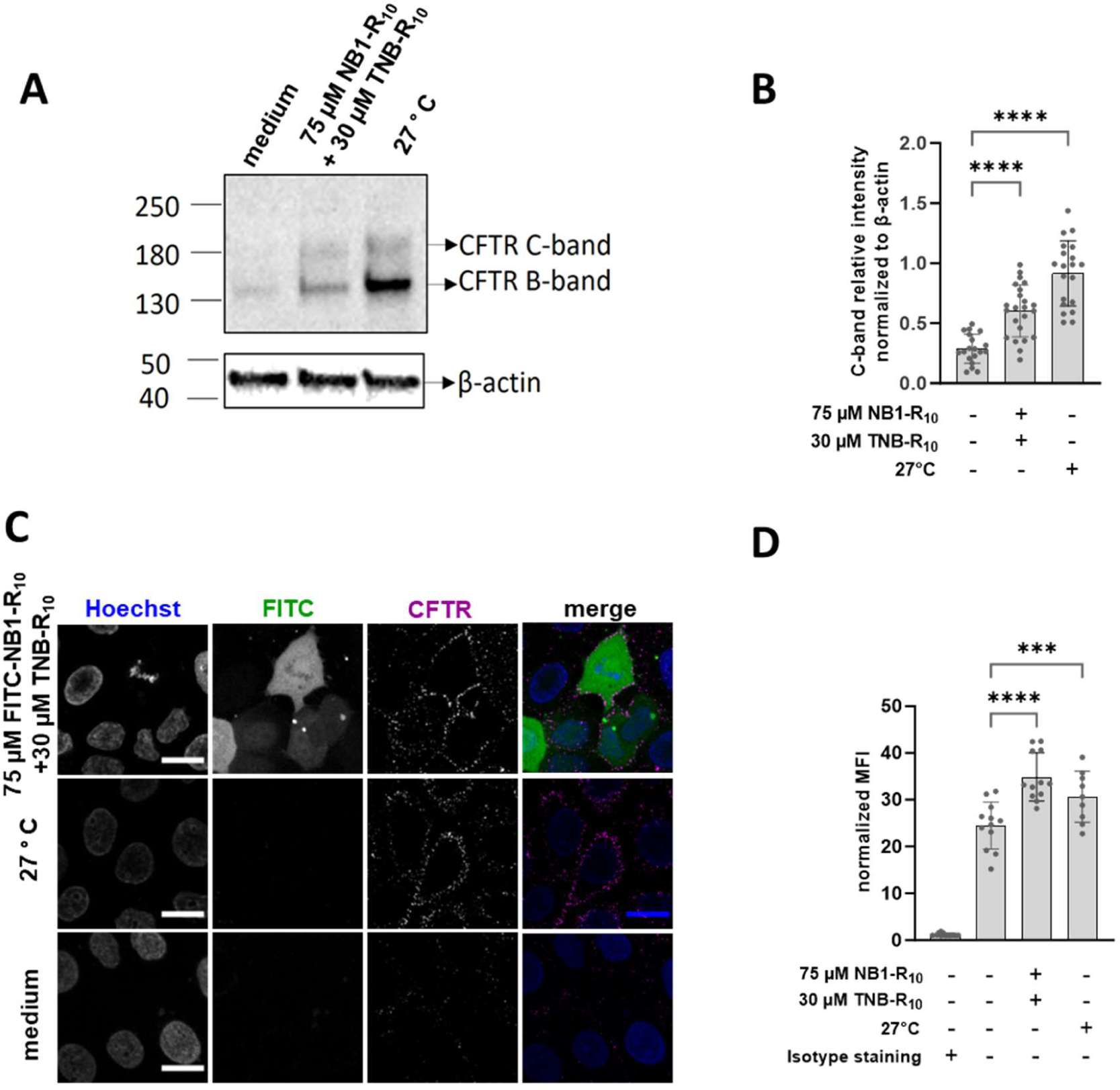
Nanobody treatment rescues F508del maturation and trafficking in CF bronchial epithelial cells. **A** Representative Western blot of medium-treated (medium), nanobody-treated (75 µM NB1-R10 + 30 µM TNB-R10), and temperature-controlled conditions (27°C) showing CFTR B-band (core glycosylated CFTR) and CFTR C-band (complex glycosylated CFTR). **B** Semi-quantification of Western blot results. Bar graph representing mean ± SEM of N=19-24 measurements of 4-6 samples with 4 technical replicates per sample and treatment group. Statistical significance is reported after comparing each group against the control (medium) with One-Way ANOVA (*** p<0.001; ****p<0.0001). **C** CFTR cell surface content determined in CFBE 41o-cells. Top row: Cells treated for 1 h with 75 µM FITC-NB1-R10/30 µM TNB-R10 in serum-free Fluorobrite DMEM and subsequently incubated for 16 h in growth medium. Middle row: Cells treated for 1 h in serum-free Fluorobrite DMEM (medium) and subsequently incubated for 16 h in growth medium. Bottom row: Cells incubated continuously in growth medium for 17 h at 27°C for temperature correction. Afterwards, cells were analyzed subsequent to CFTR antibody staining in non-permeabilized cells by live-cell confocal fluorescence microscopy (scale bar: 20 µM). **D** Semi-quantification of CFTR cell surface content by flow cytometry. For flow cytometry experiments as an antibody staining control, untreated samples were treated with an isotype control antibody (isotype staining) to exclude any unspecific staining. Data is presented as bar graph representing mean ± SEM of N= 9-12 discrete samples over 3-4 independent biological replicates, represented by dots. Statistical significance is reported after comparing each group against the control (medium) with One-Way ANOVA (* p<0.1; ***p<0.001****p<0.0001).

Next, we probed whether the maturation of F508del-CFTR upon nanobody treatment translates into trafficking of the corrected CFTR to the cell surface and results in increased CFTR cell surface levels in CFBE41o-cells. Therefore, CFBE41o-cells were incubated for 1 h with 75 µM FITC-NB1-R_10_/30 µM TNB-R_10_ in serum-free medium with a subsequent 16 h incubation in growth medium to allow for maturation and relocation of F508del-CFTR to the cell surface. Again, low temperature correction (27°C) was performed for 17 h as control^36^. To evaluate cell surface CFTR, cells were stained with a commercially available antibody against an extracellular loop peptide sequence of CFTR and subsequently either analyzed by microscopy (Fig. 2C) or semi-quantified by flow cytometry analysis (Fig. 2D). Confocal fluorescence microscopy revealed that a CFTR signal could only be observed in temperature corrected and nanobody treated CFBE41o-cells compared to CFBE41o-cells treated with medium, indicating an increase in cell surface CFTR in these conditions. For flow cytometry measurements, single, intact cells were gated and analyzed. In case of FITC-NB1-R_10_ treated samples, only the FITC-positive population (cells containing NB1) was considered in the analysis (Supplementary Fig. 2A). The APC mean fluorescence intensity (MFI) was analyzed for the appropriate populations as it is considered to correlate with CFTR levels on the cell surface in a given sample. In accordance with the microscopy results a higher normalized MFI was detected for nanobody treated cells as well as temperature corrected cells compared to the untreated control indicating a higher level of CFTR on the cell surface of these treatment groups (Fig. 2D). Accordingly, the data corroborates that the intracellular delivery of NB1-R_10_ corrects the folding and maturation of F508del-CFTR and subsequently allows its trafficking to the plasma membrane of CF bronchial epithelial cells.

To evaluate to what extent the NB1 dependent correction of maturation and folding of F508del-CFTR translates to the recovery of its chloride channel function CF bronchial epithelial cells, we monitored CFTR activity under different experimental conditions, including incubation of the unmodified CFTR-nanobody (NB1), incubation without the CPP-additive, incubation with the CPP-additive alone, incubation of an unspecific CPP-nanobody conjugate (GBP1-R_10_) and addition of a small molecule potentiator. As positive control, low temperature correction (27 °C) was performed^36^. To that end, CFBE41o-cells were cultured as confluent monolayers on semi-permeable filters and incubated apically for 24 h at aforementioned experimental conditions in serum-free medium (Fig. 3A). CFTR function was determined by measuring changes in I_sc_ in response to cAMP-dependent activation with forskolin/IBMX, and inhibition with CFTR inhibitor-172 (CFTRinh-172) as before. In a subset of experiments, the CFTR potentiator ivacaftor was added after forskolin/IBMX stimulation (Fig. 3B). Incubation with NB1, TNB-R_10_ alone, or a cell-permeable GFP-binding nanobody^17^(GBP1-R_10_) prepared in the same way as NB1, did neither increase forskolin/IBMX-induced I_sc_ nor CFTRinh-172-sensitive I_sc_ (Fig. 3C, D), confirming that restoration of CFTR function was mediated by cytosolic delivery of NB1-R_10_. We observed that incubation with 75 µM NB1-R_10_ without TNB-R_10_ additive already showed an increase in CFTR-mediated chloride current, however, not to the level measured after co-treatment with the CPP-additive. This can be expected, as cytosolic delivery of CPP-conjugated proteins can occur at higher concentrations^37^. Interestingly, in the presence of forskolin/IBMX, CFTR-mediated currents were further increased by applying the CFTR potentiator ivacaftor^38^ to NB1-R_10_-treated CFBE41o-cells. A synergistic effect of ivacaftor has been widely demonstrated when used in combination with small molecule CFTR correctors such as lumacaftor, elexacaftor, and tezacaftor^22, 23, 39^. Of note, in combination with the ivacaftor, the correction of F508del chloride channel function achieved by NB1-R_10_ treatment reached levels comparable to those of low temperature correction (Fig. 3C, D), which was previously shown to rescue F508del folding and function.^36^

**Figure 3.**
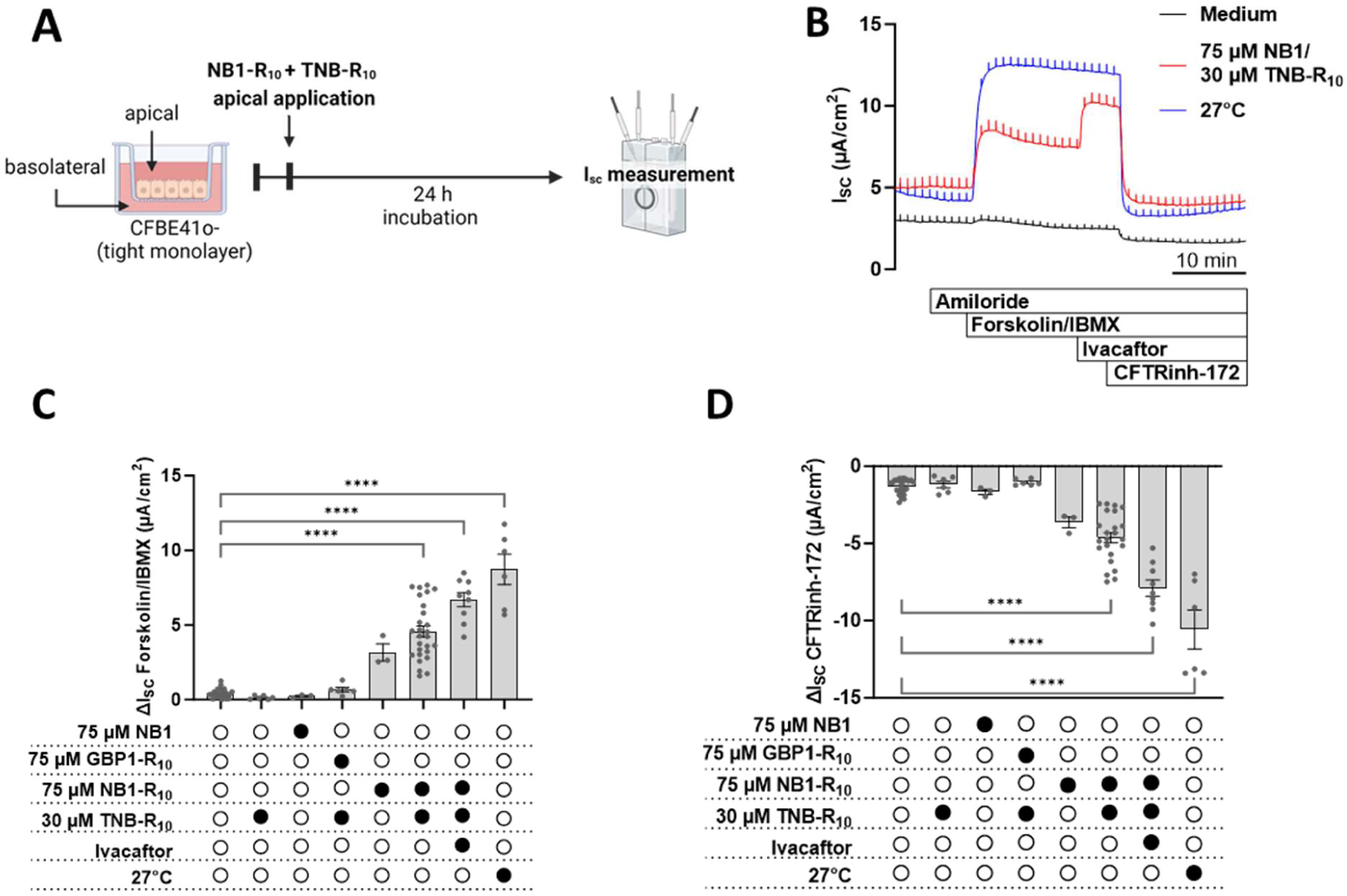
Nanobody treatment rescues F508del function in CF bronchial epithelial cells. **A** Incubation scheme for Ussing chamber experiments. **B** Representative original recordings of Isc measurements. **C-D** For the ideal conditions determined by dose response study (75 µM NB1-R10, 30 µM TNB-R10), CFTR-mediated forskolin/IBMX-induced Isc (C) and CFTRinh-172-sensitive Isc (F) were measured as indicated with and without TNB-R10 addition and in combination with CFTR potentiator ivacaftor. As additional controls, treatment with an unspecific GFP binding nanobody (GBP1-R10) and low temperature correction (27°C) were measured as indicated. Bar graphs represent mean ± SEM of N = 3 −26 samples per treatment group; each measurement is represented as dots. Statistical significance is reported for comparison of each group to vehicle by One-Way ANOVA. **** P<0.0001.

In conclusion, we were able to demonstrate that the CPP-mediated delivery of a functional CFTR-binding nanobody can modulate the fate of misfolded intracellular F508del-CFTR to restore its physiological function in a cellular model of CF. By an array of biochemical and functional assays including fluorescent cell microscopy, flow cytometry and transepithelial short-circuit current measurements, we show that the cell-permeable nanobody corrects the folding defect of the F508del protein facilitating its maturation and trafficking and thereby rescues the function of F508del-CFTR chloride channel at the apical plasma membrane.

Using the CF-causing mutation F508del in the CFTR protein as an example, this study highlights the potential of intracellular applications of extracellularly applied nanobodies for therapeutic modulation of intracellular disease mechanisms such as protein misfolding in CF and showcases the CPP-additive technology as a potential delivery tool for that purpose. Future studies will focus on translating these initial findings from a cell culture models to preclinical testing in patient-derived primary airway epithelial cells as a potential strategy to further optimize current pharmacological rescue of F508del-CFTR function towards full correction in patients with CF^20^. Beyond CF, intracellular application of nanobodies may be a promising therapeutic approach to restore protein folding and function in many other rare genetic diseases with high unmet medical need.

## Methods

### General

Solvents (DMF, CH_2_Cl_2_) were purchased from Thermo Fisher Scientific (USA). Amino acids, rink amide resin and coupling reagents were purchased from Iris Biotech (Germany). FITC HATU was purchased from Bachem (Switzerland). DIEA and TFA were purchased from Carl Roth (Germany). Salts, LB medium, antibiotics and other buffer components were purchased from Carl Roth (Germany). Mammalian cell culture media and fetal bovine serum were purchased from VWR (USA). UPLC-UV traces were obtained on a Waters H-class instrument equipped with a Quaternary Solvent Manager, a Waters autosampler and a Waters TUV detector with an Acquity UPLC-BEH C18 1.7 μm, 2.1x 50 mm RP column. The Empower 3 software (Waters) was used. Preparative HPLC of peptides was done on a Gilson PLC 2020 system using a Nucleodur C18 Htec Spum column (Macherey-Nagel, 100 A, 5 m, 250 mm x 32 mm, 30 mL/min). The following gradient was used in all purifications: A = H2O + 0.1% TFA, B = MeCN + 0.1% TFA 5% B 0-10 min, 5-50% B 10-60 min, 50-99% 60-80 min. High resolution mass spectra were measured on a Xevo G2-XS QTof (Waters) mass spectrometer coupled to an acquity UPLC system running on water and acetonitrile, both with 0.01% formic acid using the MassLynx software (V4.1, Waters). Protein spectra were deconvoluted using the MaxEnt 1 tool. For SDS-PAGE analysis proteins were mixed with 4x Laemmli buffer (Bio-Rad) with or without addition of 10 % β-mercaptoethanol (β-ME) and boiled at 95° C for 5 minutes before separation on 4–20% Mini-PROTEAN® TGX™ Precast Protein Gels (BioRad). In-gel fluorescence was imaged first, followed by Coomassie staining and imaging. Gels were imaged on a ChemiDoc XRS+ system (Bio-Rad) using the Image Lab software (Version 5.1, Bio-Rad).

### Peptide Synthesis

The TNB-R10 (ac-C(TNB)-PEG-PEG-RRRRRRRRRR, PEG: 8-amino-3,6-dioxaoctanoic acid, A) and FITC-LPETGG (Fluoresceine-LPETGG) (B) peptides were synthesized by (automated) standard fluorenylmethoxycarbonyl (Fmoc)-solid-phase peptide synthesis (SPPS) on Rink amide resin (0.05 mmol scale, 0.22 mmol/g). Arginine was incorporated with Pbf protection, cysteine was incorporated on the *N*-terminus with S-tbutyl protection. Selective deprotection of Fmoc-protected resin and Fmoc protected amino acids was achieved with 20% piperidine in DMF. Amino acid coupling was performed with 5 eq. of amino acid, 5 eq. of *O*-(1H-6-Chlorobenzotriazole-1-yl)-1,1,3,3-tetramethyluronium hexafluorophosphate (HCTU), 4 eq. of Ethyl cyanohydroxyiminoacetate (Oxyma) and 10 eq. of *N,N*-Diisopropylethylamine (DIEA) in DMF. The FITC coupling was done on the *N*-terminus after Fmoc deprotection using 2 eq. FITC (5(6)-Carboxyfluorescein, Fluka 21877) 2 eq. Hydroxybenzotriazole (HOBt) 2 eq. *N,N*′-Diisopropylcarbodiimide (DIC) overnight at RT. For TNB conjugated peptides the *N-*terminus was acetylated with DMF : acetic anhydride : DIEA (7:2:1, (v/v/v) overnight at RT. Subsequently Cys(S-tbutyl) was deprotected using 20% β-ME at RT overnight. Cysteine activation was done with Ellmans reagent overnight in Ethanol : DMF (3:1, v/v) solution. The final peptides were deprotected and cleaved off the solid support overnight (TNB-R10) or for 1 h (FITC-LPETGG) with 95% TFA, 2.5% TIS, 2.5% H_2_O. The peptide was precipitated in diethyl ether. Purification was done using HPLC with a gradient of 10 %-50 % acetonitrile (CH_3_CN) in water, both containing 0.1 % TFA over 50 min. The HPLC purification yielded a A white or B yellow green TFA salt. A: 23 mg, 0.0114 mmol, 23%; HR-MS [M+3H]^3+^ exp. 738.0032; calc.: 738.0696 B: 12 mg, 0.0129 mmol, 25%; HR-MS [M+H]^+^ exp.: 465.6764; calc.:465.6795

### NB 1 expression and purification

BL21(DE3) cells were transformed with the corresponding plasmid. A streak of colonies from a Luria Bertanni (LB)-agar plate was used to inoculate a 150 mL pre culture in Carbenicillin containing terrific broth (TB) medium and incubated overnight at 37°C under agitation. 6 L of Carbenicillin containing TB medium were supplemented with 0.1% glucose and 2 mM MgCl_2_ and inoculated with overnight with the preculture to a starting OD_600_ = 0.1. Cultures were incubated at 37°C at 180 rpm until an OD_600_ = 0.7 was reached. Cultures were then induced with 1 mM adding isopropyl ß-D-1-thiogalactopyranoside (IPTG) and further incubated overnight at 28°C, 150 rpm. Expression cultures were harvested by centrifugation (5000 xg, 10 min, 4°C) and resuspended in cold lysis buffer (0.2 M Tris pH 8.0, 0.5 M sucrose, 1 mM PMSF) corresponding to pellet weight (1:1 V/w). Solution was homogenized and stirred for 1 h at 4° C. Double the original amount of four-time diluted lysis buffer was added and further stirred for 45 min at 4°C. To remove cell debris and whole cells and clear the lysate cells were centrifuged (debris centrifugation; 25000 x*g*, 4 °C, 30 min). The supernatant was reduced for 30 min with 1-2 mM DTT.

1.5 mL Indigo beads were equilibrated with 50 mM Sodium phosphate buffer pH 7.0 containing 1 M NaCl, 10 mM imidazole. The clear lysate was incubated with Indigo beads (PureCube 100 INDIGO Ni-Agarose, Cube Biotech) for 1 h at RT. The beads were collected in a plastic column and washed with 50 mM Sodium phosphate buffer pH 7.0 containing 1 M NaCl, 10 mM imidazole and 50 mM Sodium phosphate buffer pH 6.0 containing 1 M NaCl. The nanobody was eluted with 0.05 M CH_3_COONa, 1 M NaCl, pH 4.5 into 1 M Tris-HCl pH 7.5. A second elution was performed with 0.05 M CH_3_COONa pH 4.5, 1M NaCl, 500 mM imidazole into 1 M Tris-HCl pH 7.5. The solutions were combined. TEV-cleavage was performed overnight (1:10 w/w) in dialysis against 20 mM Hepes pH 7.5, 150 mM NaCl, 10% glycerol at RT. The uncleaved nanobody, His-Tag and TEV were removed after reduction with 1-2 mM DTT using Indigo beads using storage buffer (20 mM Hepes pH 7.5, 150 mM NaCl, 10% glycerol). The nanobody was concentrated and stored at −80°C.

### GBP1 expression and purification

The GBP1 plasmid was a gift from Heinrich Leonhard^40^. GBP1 was expressed and purified as a DnaE intein and a chitin binding domain (pTXB1 vector system) fusion protein similarly to a previously published protocol^17^. Briefly, the corresponding plasmid was transformed into T7 express cells (New England Biolabs) and grown overnight at 37°C in 5 mL of LB-medium. 1 mL of this pre-culture was used to inoculate a 250 mL LB medium culture containing ampicillin. The culture was incubated at 37°C to an OD_600_=0.6. Protein expression was induced with 1 mM IPTG, and the culture was further incubated for 16 h at 18°C. Cells were harvested by centrifugation (4000xg, 15 min, 4°C). Cells were resuspended in lysis buffer ((20 mM Tris-HCl, pH 8.5, 500 mM NaCl, 1 mM EDTA, 0.1% Triton X-100, 100 µg/mL lysozyme and 25 µg/mL DNAse I), lysed by sonication (3x 2 min, 30% Amplitude), followed by debris centrifugation (25’000xg, 30 min, 4 °C). the supernatant was loaded on chitin-agarose beads, equilibrated in EPL buffer (20 mM Tris-HCl pH 8.5, 500 mM NaCl), and washed with 20 column columns of EPL buffer. The intein cleavage was performed in presence cysteine to obtain GBP1 with a c-terminal cysteine residue. Therefore, the reaction was performed on the column with 1 mM cysteine in 20 mM Tris-HCl pH 8.5, 500 mM NaCl and 100 mM sodium 2-mercaptoethanesulfonate for 16 h while shaking at room temperature. The protein was subsequently washed off the column using EPL buffer. Further purification of the reaction mixture was performed by size exclusion chromatography using a BioRad FPLC system on a Superdex 75 16/60 column in 5 mM HEPES at pH 7.5, 140 mM NaCl, 2.5 mM KCl 5 mM Glycine protein aliquots were shock-frozen and stored at −70 °C.

### Protein Conjugation

For fluorescent *N*-terminal labelling of NB1 sortase A reaction was performed. Therefore, a reaction was set up with 50 µM purified NB1. 1.1 eq. sortase A 5M, 20 eq. FITC-LPETGG peptide in sortase buffer (50 mM Tris-HCl pH 7.5, 150 mM NaCl, 0.01 mM CaCl_2_, 10% Glycerol). The reaction was performed at RT for 20 min under mild agitation. His-tagged sortase A 5M was removed via Ni-NTA. Excess peptide was by desalting in spin column using ZebaSpin 7kDa MWCO spin column (Thermo) against protein buffer.

For the CPP conjugation cysteine containing nanobodies were reduced for 20 min with 1-2 mM DTT. DTT was removed by desalting in spin column using ZebaSpin 7kDa MWCO spin column against protein buffer. 3-5 eq. of TNB-R_10_ were added directly after desalting. The solution was incubated overnight at 4° C under gentle agitation. Removal of excess TNB-R_10_ was done by desalting in spin column using ZebaSpin 7kDa MWCO spin column against protein buffer. Purification was done via desalting in Spin column.

### Mammalian cell culture

All cell lines were grown at 37° C, 5% CO_2_ in a humidified atmosphere. All cell lines used with their corresponding growth medium can be found in supplementary table 1. Cells were split at 70%-90% confluency and used in passages 4-15. HeLa CCL-2 cells and HEK cells were grown in DMEM 4.5 g/L Glucose + 10% fetal calf serum (FCS), A549 cells were grown in DMEM/Ham’s F-12 + 10% FCS, F508del-CFTR-overexpressing cystic fibrosis bronchial epithelial cells (CFBE41o-) were generously provided by Dr. Eric J. Sorscher (University of Alabama, Birmingham, USA)^41^. Cells were cultured in minimum essential medium enriched with 10% fetal bovine serum, 10 mg/mL glutamine, 100 U/ml penicillin, 100 µM/ml streptomycin and 4 µg/ml puromycin in a humidified incubator at 37 °C with 5% CO_2_.

### Cellular Uptake experiments

For microscopy experiments 50’000 cells per well were seeded in an 8-well ibidi glass bottom plate. Cells were incubated for 24 h at 37°C with 5% CO_2_ to settle. Cells were washed once with FluoroBrite DMEM without FCS or glutamine before addition of protein samples (FITC-NB1-_R10_) and CPP-additive (TNB-R_10_) in FluoroBrite DMEM without FCS or glutamine. Cells were incubated for 1 h at 37°C, 5% CO_2_. The cells were then washed three times with FluoroBrite DMEM with 10% FCS. For 1 h uptake experiments cells were counterstained with 10 µg/mL Hoechst 33342 in FluoroBrite DMEM with 10% FCS for 10 min and imaged in FluoroBrite DMEM with 10% FCS. For 24 h uptake experiments cells were further incubated for 23 h in growth medium prior to counterstaining with Hoechst 33342 as described before and imaged in FluoroBrite DMEM with 10% FCS. Imaging was done using the Zeiss LSM780 confocal microscope with a 63x, 1.4-NA Plan-Apochromat lens at RT. Image analysis and processing was performed with FIJI software^42^.

### Cell-Viability assay

For cell viability measurements using Mitochondrial dehydrogenase (WST-1) assay 15’000 cells per well were seeded in a 96-well plate and incubated for 24 h at 37°C, 5% CO2 to settle. Cells were washed once with FluoroBrite DMEM without FCS or glutamine. Cells were treated with NB-R_10_ in indicated concentrations together with 10 µM TNB-R_10_ in FluoroBrite DMEM without FCS or Glutamine for 1 h at 37°C, 5% CO_2_. Cells were washed with growth medium and incubated for 24 h at 37°C, 5% CO_2_ further. 10 µL if WST1 reagent was added to each well and incubated for 90 min at 37°C, 5% CO_2_. Absorbance at 460 nm was measured using a M200Pro plate reader (TECAN).

### Culture and treatment of cystic fibrosis bronchial epithelial cells

Prior to seeding, CFBE41o-underwent trypsinization and were passaged at a ratio of 1:2 while puromycin was removed from the culture medium. The next day, cells were trypsinized and 500,000 cells were seeded onto Snapwell inserts (Corning #3407). The CFBE41o-monolayers were maintained at a liquid-liquid interface. Upon reaching transepithelial electrical resistance (TEER) values of ≥ 600 Ω*cm^2^, the cells were subjected to apical treatment with either minimum essential medium alone (vehicle control), or different concentrations of NB1-R_10_ and TNB-R_10_ additive for 24 hours. For temperature correction of F508del-CFTR, cultures were placed in a humidified incubator at 27 °C with 5% CO_2_ for 24h.

### Ussing chamber measurements

Transepithelial short-circuit (I_sc_) measurements were conducted using EasyMount Ussing chambers (Physiologic Instruments) with a chloride gradient (145 mM basolateral and 5 mM apical) using a voltage clamp setup as previously described^43^. In brief, CFBE41o-monolayers were first equilibrated for 15-20 minutes in the presence of amiloride (100 µM). To assess CFTR-mediated chloride secretion, forskolin (10 µM) and 3-isobutyl-1-methylxanthin (IBMX, 100 µM) were administered both apically and basolaterally, followed by CFTR inhibitor-172 (CFTRinh-172, 20 µM) apically. In a subset of experiments, ivacaftor (5 µM) was administered after forskolin/IBMX to assess potentiation of CFTR-mediated currents.

### Immunoblot analysis

F508del-CFBE cell lysates were prepared by scraping the cells directly from filters into 80 µL RIPA buffer (Thermo Fisher Scientific) containing 2% sodium dodecyl sulfate and 1 mM PMSF protease inhibitor (Thermo Fisher Scientific). Following a 30-minute incubation on ice with periodic vortexing, the samples were centrifuged at 18,000 x g and supernatants were collected. For Western blotting, 5 µL of lysates were mixed with Laemmli buffer and 0.05% β-mercaptoethanol at 37 °C for 30 minutes, then subjected to electrophoresis on 4-20% gradient polyacrylamide gels (Bio-Rad). Proteins were transferred to a polyvinylidene difluoride (PVDF) membrane using a Trans-Blot Turbo (Bio-Rad) semi-dry blotting system according to the manufacturer’s instructions. The PVDF membranes were blocked in 5% milk in PBS at room temperature for 1 hour. CFTR was probed using the monoclonal CFTR antibody 596 (provided by John Riordan, Ph.D., University of North Carolina at Chapel Hill, via the CF Foundation’s antibody distribution program) at a 1:500 dilution overnight at 4 °C, followed by incubation with a horseradish peroxidase (HRP)-conjugated goat polyclonal anti-mouse antibody (Dako Denmark, P0047) at a 1:5000 dilution. The β-actin loading control was probed using monoclonal β-actin antibody (Santa Cruz, sc-477778) at a 1:2000 dilution, followed by the anti-mouse antibody. The β-actin and secondary antibodies were incubated in 2.5% milk in PBS with 0.1% Tween 20 at room temperature for 1 hour. The HRP signal was detected using the Pierce ECL Western Blotting Substrate (Thermo Fisher Scientific) and imaged using the ChemiDoc MP (Bio-Rad) imaging system. FIJI software was used for densitometric analysis^42^ . To assess CFTR maturation, the relative amount of CFTR C-band protein was normalized to loading control.

### Flow Cytometry

For the flow cytometry experiments CFBE 41o-cells were seeded with 150’000 cells in a 24-well plate. Cells were incubated for 24 h at 37°C, 5% CO_2_ to settle, then treated with 75 µM FITC-NB-R10 and 30 µM TNB-R10 or growth medium for 1 h in FluoroBrite DMEM without FCS and washed after 1 hour. They were further incubated for 16 h in growth medium. For the temperature correction control, cells were incubated in growth medium at 27° C, 5% CO_2_ for 17 h. Multiple wells per experiment were treated under the same conditions. Cells were detached with accutase, Samples were prepared at 4° C. Therefore, cells were washed twice with cold blocking buffer (PBS containing 1% BSA). Cells were stained with primary antibody against an extracellular loop peptide sequence of CFTR (CFTR Monoclonal Antibody (CF3), Invitrogen, #MA1-935, 1:200), isotype control (Mouse IgM Isotype Control (11E10), eBioscience™, Invitrogen, #14-4752-82, 1:200) or medium as untreated control, for 30 min on ice. Cells were washed twice with blocking buffer and stained with secondary antibody (Goat anti-Mouse IgM (Heavy chain) Secondary Antibody, Alexa Fluor™ 647, Invitrogen, #A21238, 10 µg/mL) or medium for untreated control for 30 min on ice. Cells were washed twice with blocking buffer and once with PBS. Cells were resuspended in PBS and measured at n a LSRFortessa (BD Biosciences, USA) flow cytometer using the LSR Fortessa cell analyzer software with 10000 events per measurement. FITC and AF 647 fluorescence were measured alongside Forward scatter (FWS) and Sideward scatter (SWS). Cell fragments and multiplets were removed in the analysis by gating using the FlowJo software. Subsequently, the FITC channel in a medium treated sample was used to determine the FITC negative population. Higher fluorescence values in FITC were deemed as the FITC positive population. The gating strategy is illustrated in suppl. Fig. 4A for a nanobody and medium treated sample. For normalization of AF647 fluorescence values, mean fluorescence value of a given sample was divided by the mean fluorescence value of the isotype control of medium treated cells.

### Live cell confocal fluorescence microscopy for cell surface CFTR staining

Cells were seeded in with 15’000 cells per well were seeded in a 96-well plate (thin bottom corning) and incubated for 24 h at 37°C, 5% CO2 to settle, then treated with 75 µM FITC-NB-R10 and 30 µM TNB-R10 or growth medium for 1 h in FluoroBrite DMEM without FCS and washed after 1 hour. They were further incubated for 16 h in growth medium. For the temperature correction control, cells were incubated in growth medium at 27° C, 5% CO_2_ for 17 h. Cells were washed once with PBS and incubated in blocking buffer (PBS containing 1% BSA). Cells were stained with primary antibody (CFTR Monoclonal Antibody (CF3), Invitrogen, #MA1-935, 1:200), isotype control (Mouse IgM Isotype Control (11E10), eBioscience™, Invitrogen, #14-4752-82, 1:200) or medium as untreated control, for 30 min on ice. Cells were washed twice with PBS and stained with secondary antibody (Goat anti-Mouse IgM (Heavy chain) Secondary Antibody, Alexa Fluor™ 647, Invitrogen, #A21238, 10 µg/mL) or medium for untreated control for 30 min on ice. Cells were counterstained with 10 µg/mL Hoechst 33342 in FluoroBrite DMEM with 10% FCS for 10 min and imaged in FluoroBrite DMEM with 10% FCS. Imaging was done using the Zeiss LSM780 confocal microscope with a 63x, 1.4-NA Plan-Apochromat lens at RT. Image analysis and processing was performed with FIJI software^42^.

### Software

Microscopy pictures were processed with ImageJ including the FIJI package. Graphing and statistics were done using Graphpad Prism 10 and Biorender. Flow cytometry data was processed and analyzed using FlowJo.

### Statistics and Reproducibility

The statistical tests and values of *N* are reported in the figure legends where appropriate. Each experiment was independently repeated at least once with similar results.

## Acknowledgements

We thank Ines Kretschmer and Yasemine Kirimlioglu for peptide synthesis and protein expression. Additionally, we thank Kathrin Seidel and Marika Drescher for support with cell culture.

## Grant support

This study was supported by the German Research Foundation (CRC 1449 – project 431232613, sub-project A01, C04 and Z02 to M.A.M. and C01 to C.P.R.H. as well as RTG2473 “Bioactive Peptides” – project 392923329 to C.P.R.H.), the German Federal Ministry of Education and Research (82DZL009B1 to M.A.M.), the Muko e.V. and by Dr. Kurt Schmalz and J. Schmalz GmbH to C.P.R.H. C.G. acknowledges support by the Fond Forton, the Welbio (grant WELBIO-CR-2022 A – 09), the Association Luxembourgeoise de Lutte contre la Mucoviscidose, the Fondation Air Liquide and the FRS-FNRS.

## Author information

These authors contributed equally: Luise Franz, Tihomir Rubil

## Contribution

CG, MAM and CPRH conceived the project. LF, TR, AB, MAM, CPRH designed the project and experiments. MO designed the nanobody plasmid and supplied the initial expression and purification protocol. LF and KKH performed the expression optimization and purification. LF performed the protein functionalization, cellular uptake experiments, microscopy and flow cytometry. TR and AB performed the functional studies and Western blot experiments. LF and TR analyzed all data and wrote the manuscript. All authors edited and approved the manuscript.

## Extended Figures

**Extended Data Figure 1:**
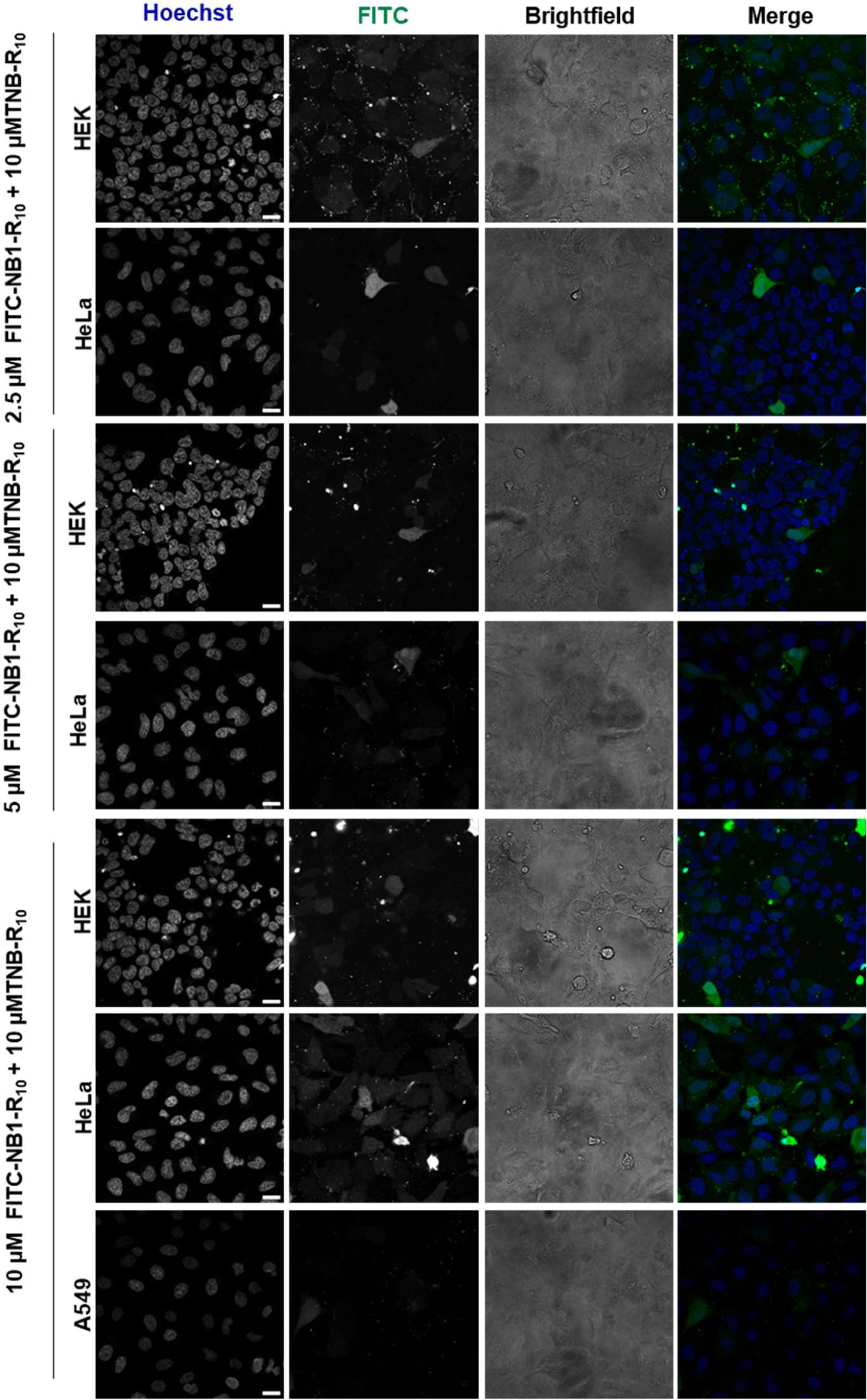
Cellular uptake of cell permeable NB1-R10 into different cell lines. HEK and A549 cells were incubated with 2.5 −10 µM FITC-NB1-R10 (FITC) and 10 µM TNB-R10 in serum free FluoroBrite DMEM for 1 hour at 37 °C, counterstained with Hoechst and imaged by live-cell confocal microscopy in FluoroBrite DMEM with 10% FCS at RT. Scale bar: 20 µM.

**Extended Data Figure 2:**
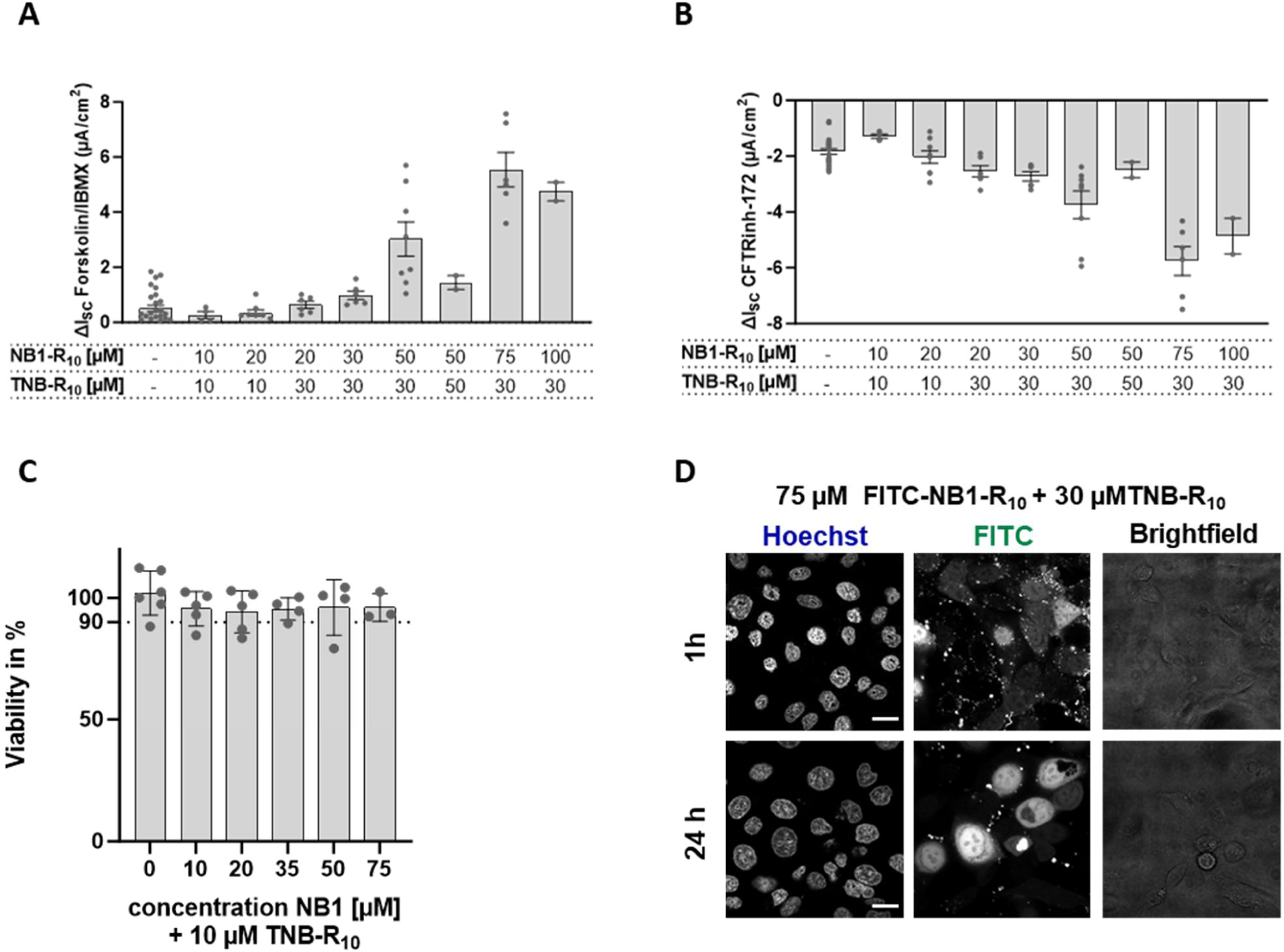
**A-B** Initial dose-response study to evaluate the effects of treatment with different concentrations of NB1-R10 (0-100 µM) in combination with CPP additive (TNB-R10, 10-50 µM) on forskolin/IBMX-induced Isc (C) and CFTRinh-172-sensitive Isc (D) Bar graphs represent mean ± SEM of N= 2-27 samples, each represented as dots. **C** Cell viability measured by WST-1 assay in CFBE 41o-cells. Absorbance at 440 nm indicates cellular metabolic activity in the processing of WST-1 to the absorbing Formazan. The treatment with 0-75 µM NB1-R10 in presence of 10 µM TNB-R10 for 1 h and a subsequent 24 h incubation in growth medium had no effect on cell viability. Data presented as bars representing the Mean ± SEM of *n*=2 biological replicates with 1-3 discrete samples per replicate represented as dots. **D** Live-cell confocal microscopy images of cell permeable nanobody into CFBE41o-cells with 75 µM FITC-NB1-R10/30 µM TNB-R10 directly after 1 h of incubation in serum-free FluoroBrite DMEM (1h) or after subsequent 24 h of incubation in growth medium (24h). Cells were counterstained with Hoechst prior to imaging at RT. Scale bar: 20 µM.

## Notes

### Competing Interest Statement

The authors have declared no competing interest.

