## Supplemental Information for "A Cell-Permeable Nanobody to Restore F508del Cystic Fibrosis Transmembrane Conductance Regulator Activity"

**Supplementary Information**

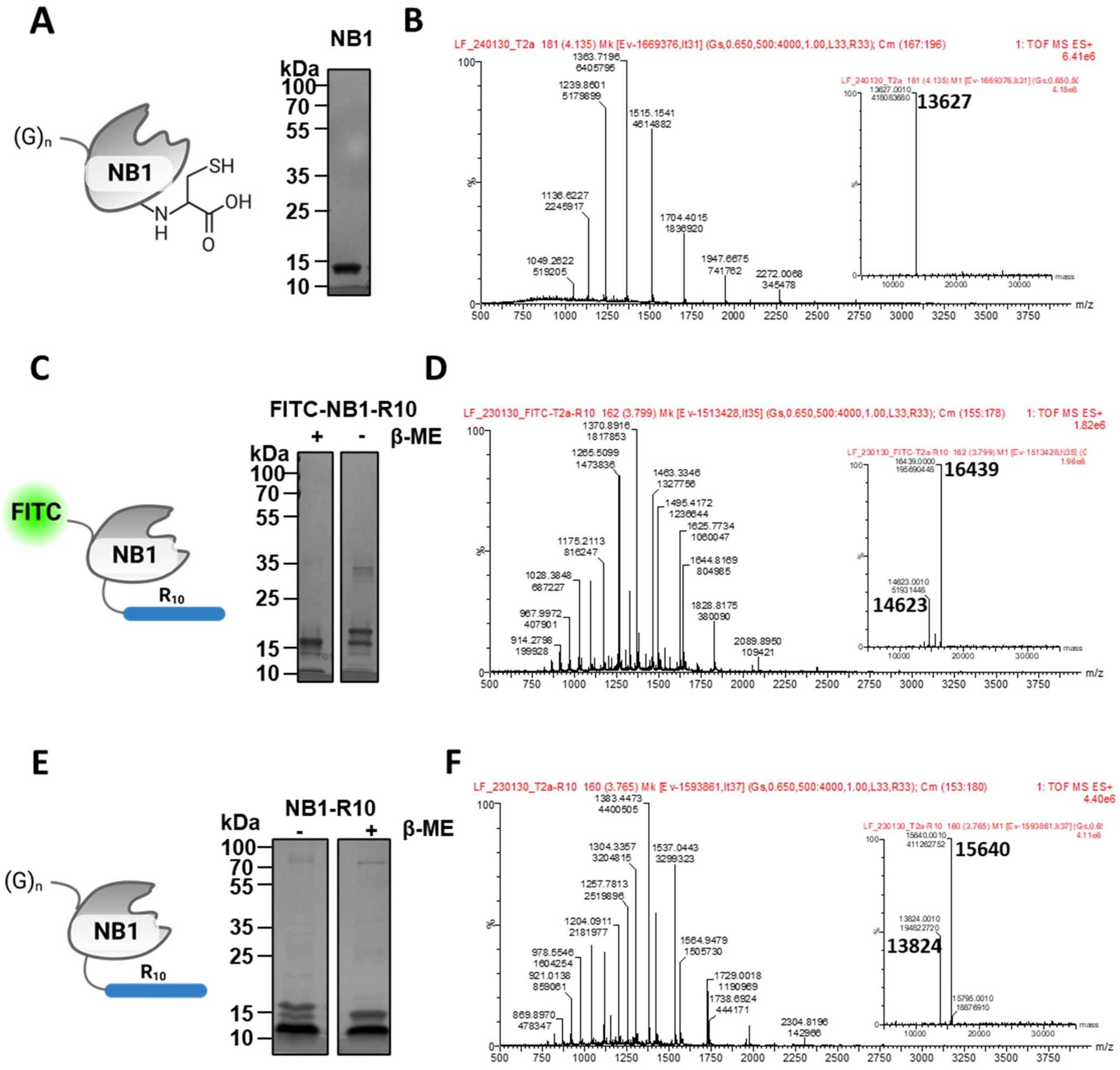


**Supplementary Figure 1**: Characterisation of purified nanobody conjugates. **A** SDS PAGE gel showing purified NB1 in 1x Laemmli buffer after purification. **B** HRMS spectrum of NB1 [M+H]$^+$ calc. 13627Da. **C** SDS-PAGE gel of FITC-NB1-R10 in 1x Laemmli buffer with (+) or without (-) 2.5% β-mercaptoethanol (β-ME) showing two bands without reducing conditions indicating the succesfull conjugation of CPP via disulfide. **D** HRMS spectrum of FITC-NB1-R10 [M+H]$^+$ calc. 16439 Da, FITC-NB1 [M+H]$^+$ calc. 14632 Da. **E** SDS-PAGE gel of NB1-R10 in 1x Laemmli buffer with (+) or without (-) 2.5% β-ME showing two bands without reducing conditions indicating the succesfull conjugation of CPP via disulfide. **F** HRMS spectrum of NB1-R10 [M+H]$^+$ calc. 15640 Da.

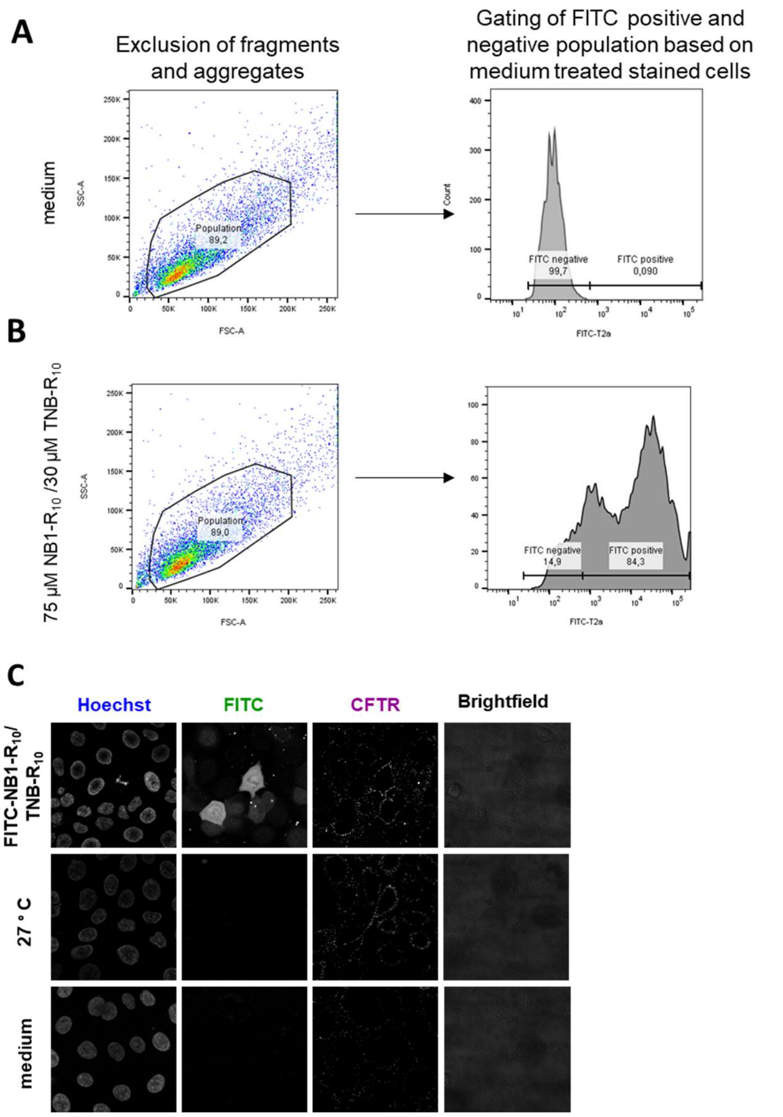


**Supplementary Figure 2**: Investigation of cell surface CFTR **A** Gating example for semi-quantitative analysis of CFTR on the cell surface by flow cytometry shown for a stained untreated sample (medium). To select the viable population of a sample sideward scatter (SSC-A) and forward scatter (FSC-A) were used. The gate was set to exclude fragments and aggregates. For the nanobody treated sample only the nanobody containing, FITC positive population, was considered to determine the mean APC fluorescence representing the stained cell surface CFTR content. The gate for FITC negative population was set using the untreated (medium) sample. The population with higher fluorescence than the FITC negative population was gated as FITC positive population as indicated. **B** application of gates to an exemplary measurement of a stained nanobody treated sample (75 µM NB1-$R_{10}$ + 30 µM TNB-$R_{10}$) **C** full set of live-cell confocal microscopy images of cell surface CFTR staining (Fig. 3D). CFBE 41o- cells were treated for 1 h with 75 µM NB-$R_{10}$ and 30 µM TNB-$R_{10}$ in serum-free FluoroBrite DMEM, serum-free FluoroBrite DMEM (medium) and subsequently incubated for 16 h in growth medium or incubated in growth medium for 16 h at 27°C as temperature control (27°C)
